# Bioinformatic Analysis Reveals That Some Mutations May Affect On Both Spike Structure Damage and Ligand Binding Site

**DOI:** 10.1101/2020.08.10.244632

**Authors:** Emre Aktas

**Author notes:** Correspondence with Emre Aktaş Afyon Kocatepe University, Faculty of Art and Science, Department of Moleculer Biology and Genetics, Bioinformatic Section, ANS Campus 03200 Afyonkarahisar, Turkey.

## Abstract

There are some mutations are known related to SARS-CoV-2. Together with these mutations known, I tried to show other newly mutations regionally. According to my results which 4326 whole sequences are used, I found that some mutations occur only in a certain region, while some other mutations are found in each regions. Especially in Asia, more than one mutation(three different mutations are found in QLA46612 isolated from South Korea) was seen in the same sequence. Although I detected a huge number of mutations (more than seventy in Asia) by regions, some of them were predicted that damage spike’s protein structure by using bioinformatic tools.The predicted results are G75V(isolated from North America), T95I(isolated from South Korea), G143V(isolated from North America), M177I(isolated Asia), L293M(isolated from Asia), P295H(isolated from Asia), T393P(isolated from Europe), P507S(isolated from Asia), D614G(isolated from all regions) respectively. Also, in this study, I tried to show how possible binding sites of ligands change if the spike protein structure is damaged and whether more than one mutation affects ligand binding was estimated using bioinformatics tools. Interestingly, mutations that predicted to damage the structure do not affect ligand binding sites, whereas ligands’ binding sites were affected in those with multiple mutations.Focusing on mutations may opens up the window to exploit new future therapeutic targets.

## 1. Introduction

In two decades, mankind have come acrossed at least one lethal outbreaks caused from betacoronaviruses [1–5]. The first was Severe Acute Respiratory Syndrome Coronavirus (SARS-CoV) in 2002, which infected over 8,000 people and nearly 800 people were died [6]. In 2012, Middle East Respiratory Syndrome, MERS-CoV was followed and it was resulted in with 2,294 cases [3]. Last one, SARS-CoV-2 is the cause of the severe respiratory disease COVID-19 [7]. The first reported case was in China by the end of December 2019 [8], and triggered an epidemic that quickly spread whole world and resulted in a pandemic[5]. It is called by World Health Organization (WHO) situation report reads: over 11 million confirmed cases, and over 539 000 deaths (WHO Situation Report Number 170, July 8).

Having information about viral mutations (COVID-19) is very important for insights for assessing viral drug resistance, immune escape and pathogenesis related mechanisms, moreover; it may play a vital role for designing new vaccines, antiviral drugs and diagnostic assays. However, mutagenic process is so complex and many reasons play a role in this process such as;replicate the nucleic acids, influenced by few or no proofreading capability and/or post-replicative nucleic acid repair, host enzymes, spontaneous nucleic acid damages due to physical and chemical mutagens, recombination events and also particular genetic elements. [7, 8]. In addition, some combinations factors are thought that make COVID-19 so dangerous. They might be that humanity have no direct immunological experience with SARS-COV-2, affecting us prone to infection and some other diseases. It is quite high transmissible from man to man; and it has a very big mortality rate. Commonly range is between 2.2-3.9 and range of deaths per confirmed cases is 0.5-15% [8, 9] (Mortality Analyses, John Hopkins University of Medicine). COVID-19 which rapid globally spread may provide the virus with plentiful opportunity for natural selection to act on mutations. It might be thought like case of influenza(where mutations slowly accumulate in the hemagglutinin protein during a flu season), and there is a complex interplay between mutations that can confer immune resistance to the virus, and the fitness landscape of the particular variant in which they arise [5]. The SARS-CoV-2, which mutation rate rate remarkable high and many variation have already characterized, has shown to have gone through certain mutations both in its structural and non structural proteins within several months while spreading throughout the world [8,10]

I focused my study on both determining some mutations that occurred based on the regions and evaluating whether this new mutation had an effect on the shapes of spike proteins. Besides I tried to predict that how mutations affect on ligand site. Characterization of these detected variants may give a new way for making new vaccine design, treatments and diagnostic approach.

## 2. MATERIAL METHOD

### 2.1. Dataset construction

Almost 4526 whole sequences, which are belongs to surface glycoprotein, of COVID-19(taxid:2697049) isolated from humans have been downloaded from NCBI Virus website based on their geographic region (www.ncbi.nlm.nih.gov/labs/virus).

### 2.2. Nucleotide substitution analysis

The dataset has been aligned by using MEGAX(align by MUSCLE) program. Geographical regions were evaluated separately one by one[11].

### 2.3. Predicted protein structure

Phyre2 is a suite of tools available on the web to predict and analyze protein structure, function and mutations. All of predicted structure are obtained by this tool[12](www.sbg.bio.ic.ac.uk/phyre2/html).

### 2.4. Predicted structure models

MISSENSE3D online tool is used to predicted structure of my missense variant to compare with normal structure[13].

### 2.5. Predicted phylogenetic clusters and genotypes

Genome Detective Coronavirus Typing Tool are used. This application allows us identify phylogenetic clusters and genotypes from assembled genomes in FASTA format[14].

### 2.6. Prediction of Ligand Site

3DLigandStie method was used to do an automated for the prediction of ligand binding sites[15].

## 3. RESULTS and DISCUSSION

### 3.1. Finding mutations based on regions

After downloading whole spike sequences, I performed all different regions seperately. On Africa region, the most common mutations are Q667H (5 mutations), D614G (3 mutations), R408I (2 mutations) others (1 mutations) respectively in my 84 whole sequences and there are found eight different mutations. They are shown on table 1. Interestingly, one of them QJX45344 isolated from Tunisia has two mutations A288T, Q314R respectively. The predicted structure damage for both do not affect structure damage of spike based on resulting MISSENSE3D online tool. In this area only when D614G mutation occur, the predicted structure has damage. For others, there is no any predicted structure damage results based on their sequences after using MISSENSE3D online tool.

**Table 1.**
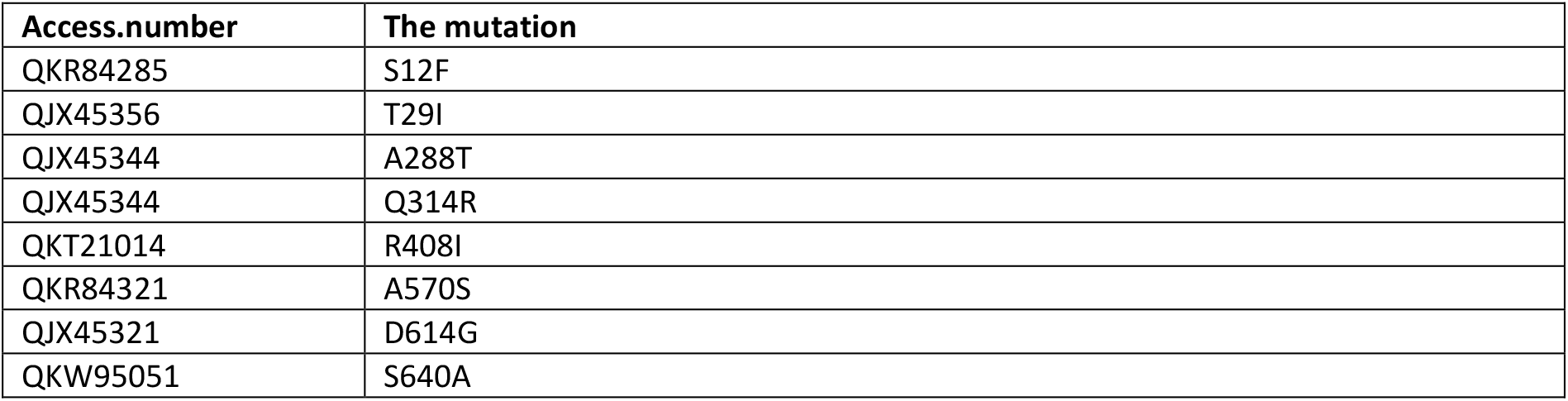
Eight different mutataions types are shown based on whole sequences of surface glycoprotein (Africa) from NCBI by using MEGAX manually.

| Access.number | The mutation |
| --- | --- |
| QKR84285 | S12F |
| QJX45356 | T29I |
| QJX45344 | A288T |
| QJX45344 | Q314R |
| QKT21014 | R408I |
| QKR84321 | A570S |
| QJX45321 | D614G |
| QKW95051 | S640A |

According to 347 whole sequences of spike protein, the most common mutations are D614G (39 mutations), H49Y(3 mutations), Y453F (8 mutations), G261D(6 mutations), A845S (4 mutations), T676I(2 mutations), S254F(2 mutations), I197V(2 mutations) respectively and others have only one mutation are shown on the table 2. It is clear that same mutation can occur different position. For example, Alanine can turn into Serine at two different positions such as;A845S, A892S. Besides, Threonine(T) can change into Isoleucine in three different positions(T22I, T240I, T676I). Only two predicted structure damage are estimated, they are T393P and D614G.

**Table 2.** Thirty five different types of mutations are shown only for Europe by helping MEGAX program.

| Access.number | The mutation | Access.number | The mutation |
| --- | --- | --- | --- |
| QKM76366 | T22I | QJS39507 | N501T |
| QJT72134 | L5F | QJT73034 | T553N |
| QHU79173 | H49Y | QJC19455 | K558R |
| QJD23141 | Q115R | QJT72470 | T572I |
| QJT72086 | M153I | QJT72278 | L611F |
| QJT72350 | L176I | QKM76846 | D614G |
| QJS53410 | N188D | QJT72614 | T676I |
| QJS53494 | I197V | QJZ28203 | M740I |
| QKJ68364 | V213L | QJS54286 | G769V |
| QJT73010 | T240I | QJS53386 | Y789D |
| QKM76906 | S254F | QIC53204 | F797C |
| QJS39543 | G261D | QJT72710 | A845S |
| QJS39627 | V367F | QJS53578 | A892S |
| QJT72806 | V382E | QJT72242 | A1020V |
| QJT72386 | C379F | QJS53506 | H1101Y |
| QJS54106 | T393P | QJS53398 | V1122L |
| QJS39603 | Y453F | QJZ28203 | D1260N |
| QJS39567 | F486L |  |  |

For 760 whole sequences from Ocenia and South America, the most finding mutations are G1124V (25 mutations), D614G (20 mutations), other different mutations tend to increase such as; S50L(10 mutations), A262T(11mutations), L5F(5mutations), D138H(3mutations), S221L(3 mutations), G485R(3 mutations). All finding mutation are shown on the table 3. Like Europe and Africa, there are same mutations occurred different position such as; T29I, T76I, T791I. Besides QKV37632 isolated whole sequences of spike protein has two mutations. They are T29I and S704L. Like all regions, when D614G occur, structure’s damage is predicted by tool.

**Table 3.** Thirty one different types of mutations are shown. 760 whole sequences from Ocenia and South America. (20 of them are belongs to South America) of spike protein were used.

| Access.Number | The mutation | Access.Number | The mutation |
| --- | --- | --- | --- |
| QJR90681 | L5F | QHR84449 | D614G |
| QKV37632 | T29I | QJR87501 | P621S |
| QKV38004 | H49Y | QJR93417 | A626V |
| QJR87081 | S50L | QKR84925 | Q675H |
| QJR88113 | T76I | QJR87477 | Q701H |
| QJR92637 | I128F | QKV37632 | S704L |
| QJR93237 | D138H | QJR85593 | M731I |
| QJR93801 | L176F | QJR88113 | T791I |
| QJR89217 | S221L | QKV38208 | P812S |
| QHR84449 | S247R | QJR87261 | A846V |
| QJR87129 | W258L | QJR93861 | D936Y |
| QJR87465 | A262T | QJR88221 | P1079S |
| QJR86937 | I468T | QKV37548 | G1124V |
| QKR86245 | G485R | QJR85701 | D1163G |
| QKR85081 | H519Q | QJR85833 | D1260N |
| QJR85965 | P561L |  |  |

North America shown on the table 4 has a quite high D614G mutatition rate based on my 2700 complete sequences of only spike proteins. I determined over 255 mutations for D614G. Based on my samples, some other mutations are seen such as; L5F (19 mutations), D138H (18 mutations), E554D ((13 mutations), P631L (10 mutations) respectively. Besides, two different mutations are found at the same position too. An example is QKG89654(A845D) and QKV35819 (A845V)have the same position but different mutation. The other example is QKG91034(Q836P) and QKG81751(Q836L). These two example may be a proof that some position more vulnarable to change into other mutations. For both, there was no predicted structure damage according to MISSENSE3D online tool. Like Europe and Africa Threonine(T) change into Isoleucine(I) in three different positions. Besides I did not find any predicted structure damage result.

**Table 4.**
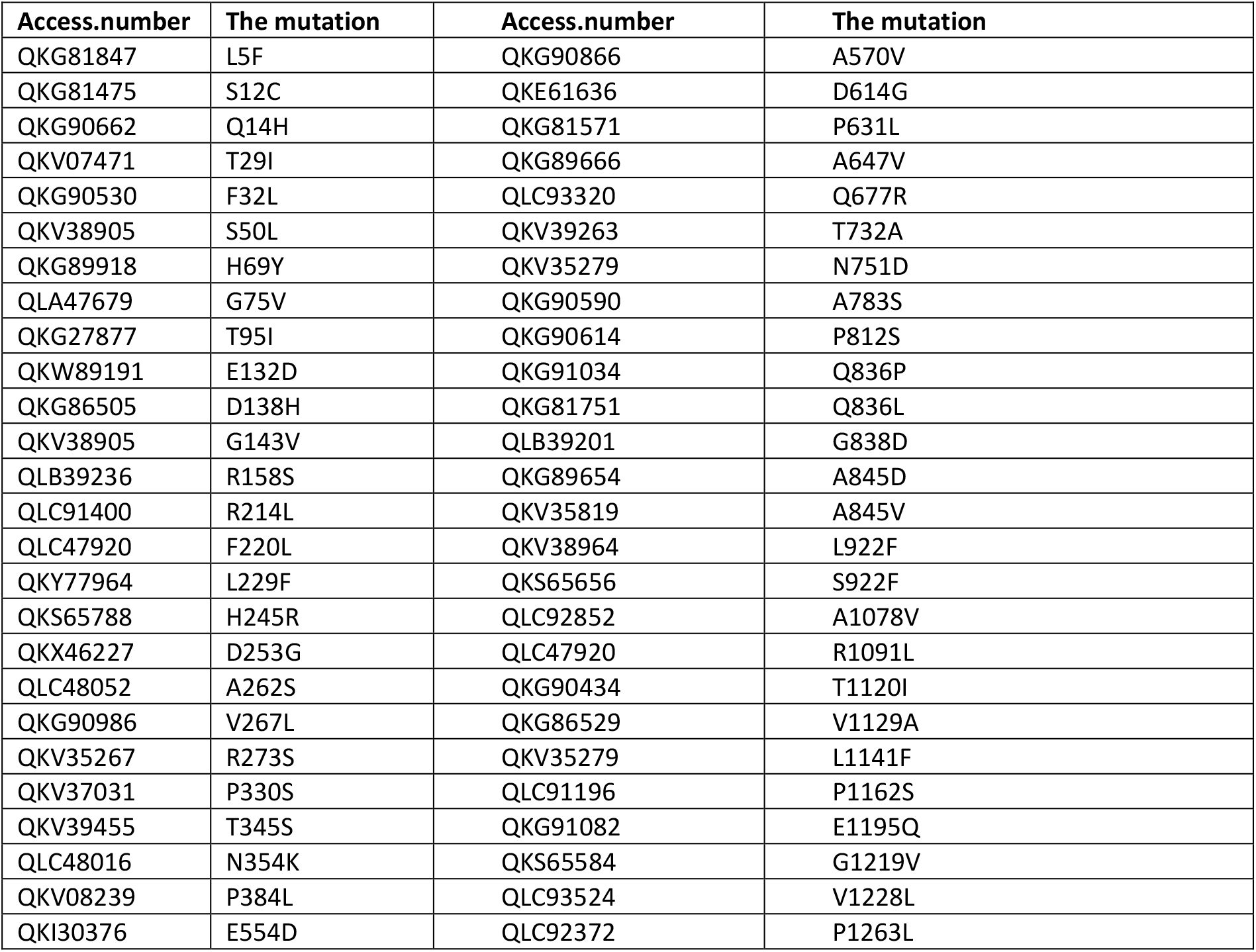
Fifty two different mutations are shown based on whole seguences (2500 sequences)of spike proteins in the North America.

Even number of sequences used are not so high according to Europe and North America, many mutations are found by using MEGAX program for Asia. It is the region where the most mutation types are seen, they are shown on the table 5. Like nearly all regions, D614G are the most variant, nearly 240 isolated samples are found. In addition, some mutations more than three L54F (40 mutations), R78M (15 mutations), V367F(5 mutations), A829T(10 mutations), H1083Q (4 mutations), T791I(12 mutations), Q677H(4 mutations), E583D(15 mutations), T572I(10 mutations), L8V(4 mutations) are found even some of other have one or two mutations. All mutations are shown on table 5. Surprisingly QLA46612 isolated from South Korea has three different mutations which are L54F, F86S, T95I and QKY60177 isolated from India has four mutations that are Q506H, P507S, Y508N, K786N. Interestingly, when all of this mutation occur, no any predict results about structure damage by using MISSENSE3D online tool. Some mutations were found more than one. One can see on the table 5 are Threonine(T) change into Isoleucine (I) occur different position such as; T22I, T76I, T95I, T572I, T791I, T827I and Glutamine (Q) turn into Histidine(H)(QLA10116, QKW92184). I found only mutations belonging to this region. Some of them areC1243F, Q1201K, K1191N, D1153Y, P507S etc. Another example of three mutation occur at the same isolated sequences is QKY60177. It has Q506H, Y508N, P507S respectively. Like QLA46612 isolated from South Korea, QJD23249 isolated from Wilayah Persekutuan Malaysia has four mutations which are L293M, D294I, P295H, H519Q. I conducted all mutations (for QJD23249) one by one, however, I did not find any predicted structure damages. These both results do not affect on structure damage according to my results after resulting of bioinformatic process.

**Table 5.** Seventy six mutations are shown based on whole seguences (635 sequences)of spike proteins in the Asia region. Only one of isolated accession number are used even many are found.

| Access.number | The mutation | Access.number | The mutation |
| --- | --- | --- | --- |
| QJX44586 | F2L | QJD23249 | H519Q |
| QIT07011 | L8V | QIU81885 | A570V |
| QJX44430 | S13I | QJT43608 | T572I |
| QKO25614 | Q14H | QKJ68545 | D574Y |
| QJQ84843 | T22I | QJR84537 | E583D |
| QIA20044 | Y28N | QKJ68497 | Q613H |
| QKO25770 | H49Y | QIT06999 | D614G |
| QIU80913 | S50L | QIU81873 | A653V |
| QKO25770 | T76I | QKT20894 | H655Y |
| QLA46612 | L54F | QKW92184 | Q675H |
| QJY40517 | R78M | QLA10116 | Q677H |
| QLA46612 | F86S | QKV49386 | R682Q |
| QLA46612 | T95I | QKN61217 | R682W |
| QKO25758 | D138H | QKU37093 | A684V |
| QKE61684 | N148Y | QJX44634 | A706S |
| QKV27551 | W152 | QJD47800 | R765L |
| QKQ30162 | M153I | QIZ16509 | V772I |
| QJT43452 | E156D | QJD20632 | T791I |
| QJY40469 | S162I | QKY60177 | K786N |
| QKJ68737 | Q173H | QKY65277 | K795Q |
| QJW00291 | M177I | QKO00486 | P809S |
| QLA09870 | K188N | QJQ84831 | A829T |
| QKO25794 | N211Y | QJT43584 | T827I |
| QHZ00379 | S221W | QJX44466 | A879S |
| QLA10140 | W258L | QJD47718 | S884F |
| QKY60121 | A262S | QJT43572 | A892V |
| QKX47933 | G261R | QIA98583 | A930V |
| QJC19491 | Q271R | QKK12815 | S939Y |
| QJD23249 | L293M | QKF95522 | Q1002E |
| QJD23249 | D294I | QJY40517 | H1083Q |
| QJD23249 | P295H | QKI31226 | F1109L |
| QKV49386 | V367F | QKO25782 | V1104L |
| QJX44562 | E471Q | QJR84369 | K1181R |
| QKY60177 | Q506H | QKJ68545 | D1153Y |
| QKY60177 | Y508N | QKO25674 | K1191N |
| QKY60177 | P507S | QKJ68605 | Q1201K |
| QKY60189 | P507H | QJR84429 | C1243F |

### 3.2. Predicted Reasons Of Structures Damages

All missense mutation are used to predict structure damage and results are shown on the figure 1. Predicted structure damage’s reason of D614G found for all regions is substitution replaces glycine originally located in a bend curvature in this area (Fig. 1A). T393P isolated from Europe substitution introduces a buried proline and it triggers disallowed phi/psi alert. The phi/psi angles are in favored region for wild-type residue but outlier region for mutant residue (Fig. 1B). The predicted reason for this M177I isolated from Asia is that substitution results in a change between buried and exposed state of the target variant residue. MET is buried (RSA 1.0%) and ARG is exposed (RSA 16.9%)[(RSA < 9% for buried and the difference between RSA has to be at least 5%. (Fig. 1C). This substitution(P507S isolated from Asia) replaces a buried uncharged residue (PRO, RSA 0.0%) with a charged residue (HIS) (Fig. 1D). Substitution (P295H isolated from Asia) replaces a buried uncharged residue (PRO, RSA 0.7%) with a charged residue (HIS) and leads to the expansion of cavity volume by 142.128 Å^3 (Fig. 1E). Substitution (L293M) results in a change between buried and exposed state of the target variant residue. LEU is buried (RSA 2.4%) and MET is exposed (RSA 13.2%)(Criterion: The substitution results in a change between buried and exposed state of the target variant residue. (RSA < 9% for buried and the difference between RSA has to be at least 5%.) (Fig. 1F). Substitution G75V isolated from North America) replaces a buried GLY residue (RSA 3.5%) with a buried VAL residue (RSA 0.0%) (Fig. 1G). This (G143V)substitution triggers disallowed phi/psi alert. The phi/psi angles are in allowed region for wild-type residue but outlier region for mutant residue and it replaces glycine originally located in a bend curvature (Fig. 1H). The substitution(T95I isolated from both Asia and North America) disrupts all side-chain / side-chain H-bond(s) and/or side-chain / main-chain bond(s) H-bonds formed by a buried THR residue (RSA 0.0%) (Fig. 1I). The phylogenetic tree of my mutations are shown on the figure 2 by using bioinformatic tools. They tend to close both bat SARS CoV and outgroup according to Genome Detective Coronavirus Typing Tool. Interestingly, mutations that damage the structure do not affect ligand binding sites (Figure 3), whereas ligands’ binding sites were affected in those with multiple mutations (Figure 4). The result of all mutations detected to be affected by the structure was the same and is shown in figure 3. For example, the same source structure (2dd8_S, 2ajf_E pdb) was taken for the structure predicted for all ligand binding. Besides, all amino acids were same (Figure 3).

**Figure 1.**
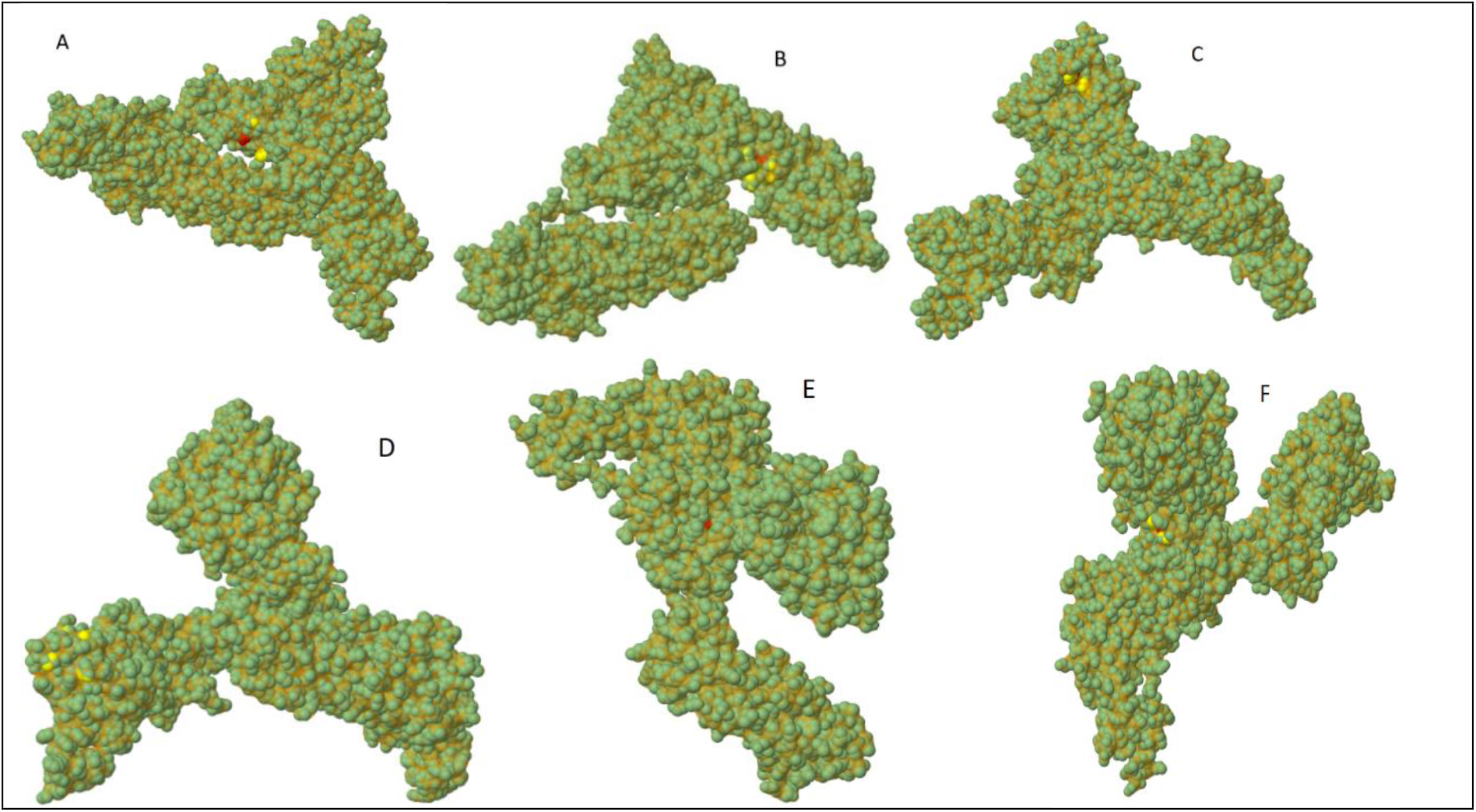
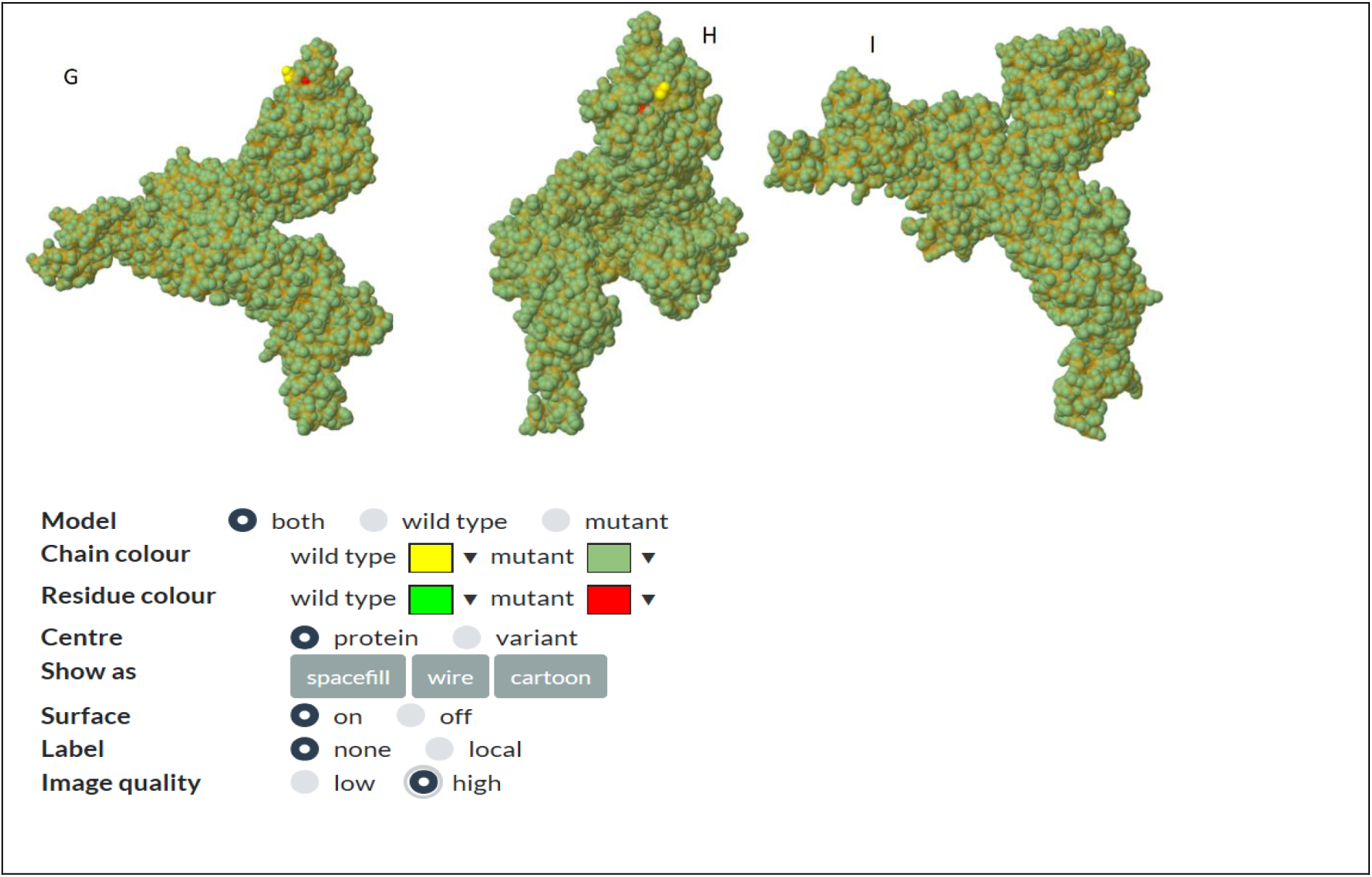
shows which mutations disrupt the structure of the spike protein. And both structure are given and it illustrates by color. While yellow color shows wild type’s chain color, dark green shows mutant chain color. Light green demonstrate is color of wild type residue color and red one shows mutant residue color. The reason why the light green color does not appear is that it remains inside the shape.

**Figure 2.**
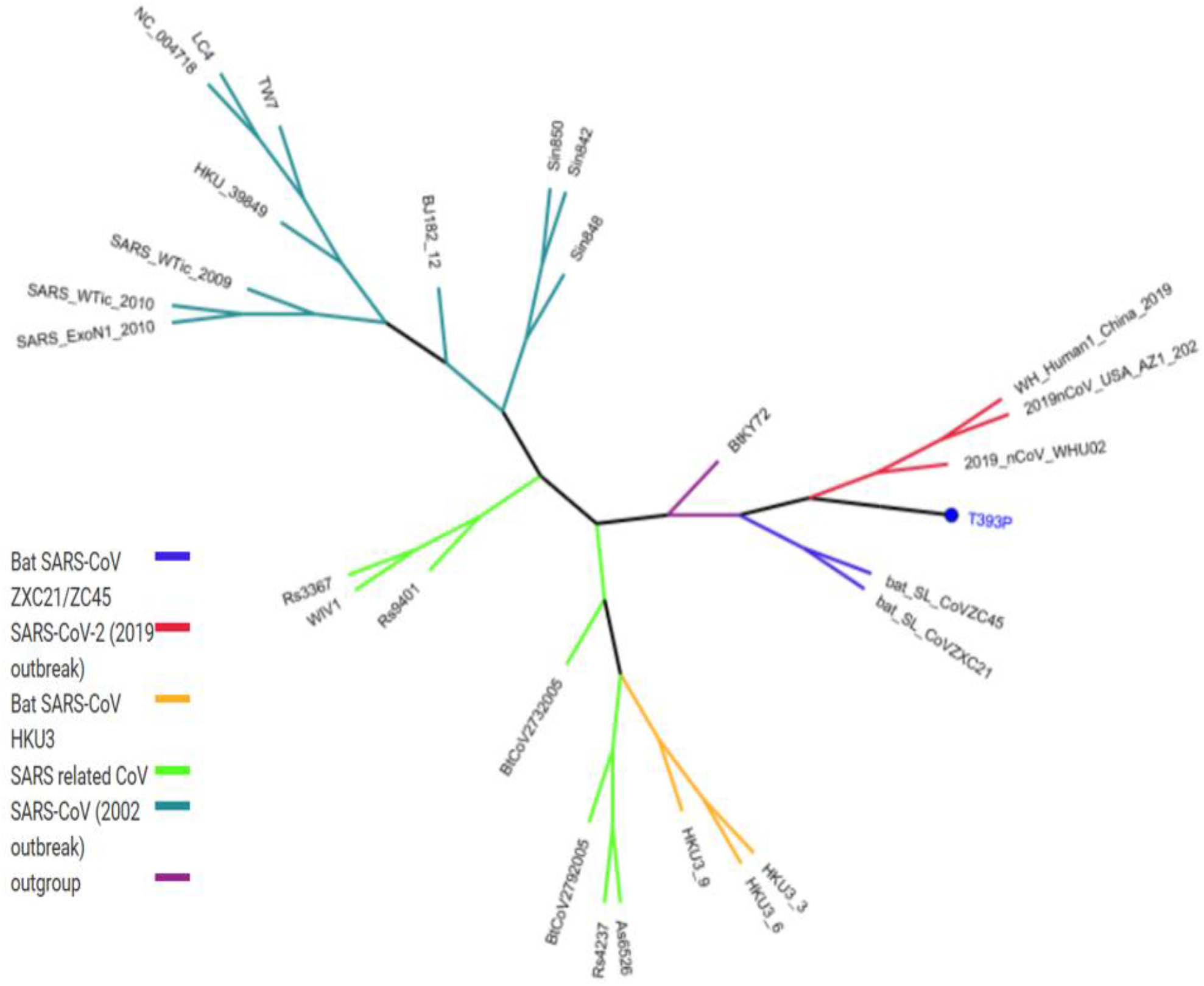
show phylogenetic tree of one mutation which are predicted to play a role in structure damage by using Genome Detective Coronavirus Typing Tool. All mutations have same location.

**Figure 3.**
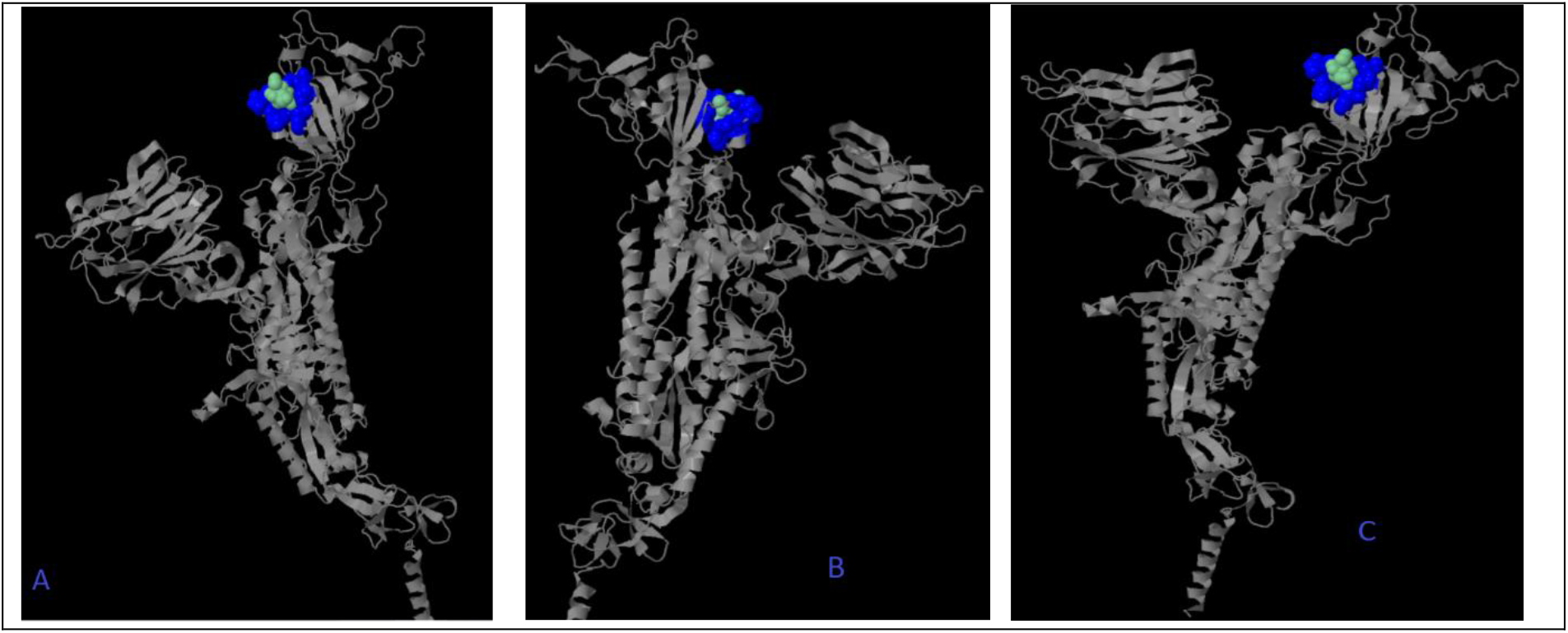
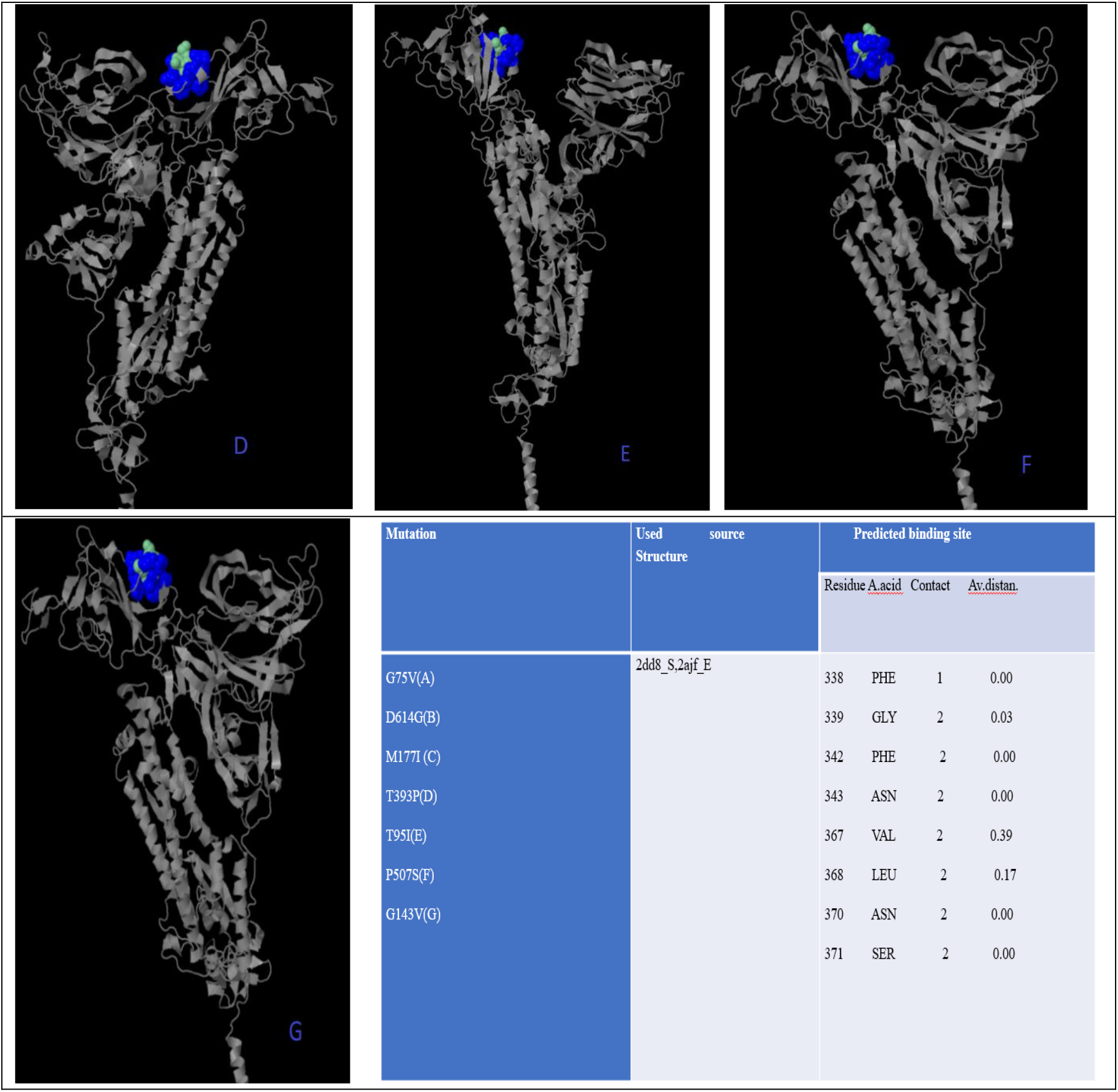
is about mutations that damage the structure do not affect ligand binding sites. Because all results(predicted ligand binding sites) were the same even their structure are different from each other. While blue color represents predicted residue, cyan represents heterogens based on 3DLigandStie method.

**Figure 4.**
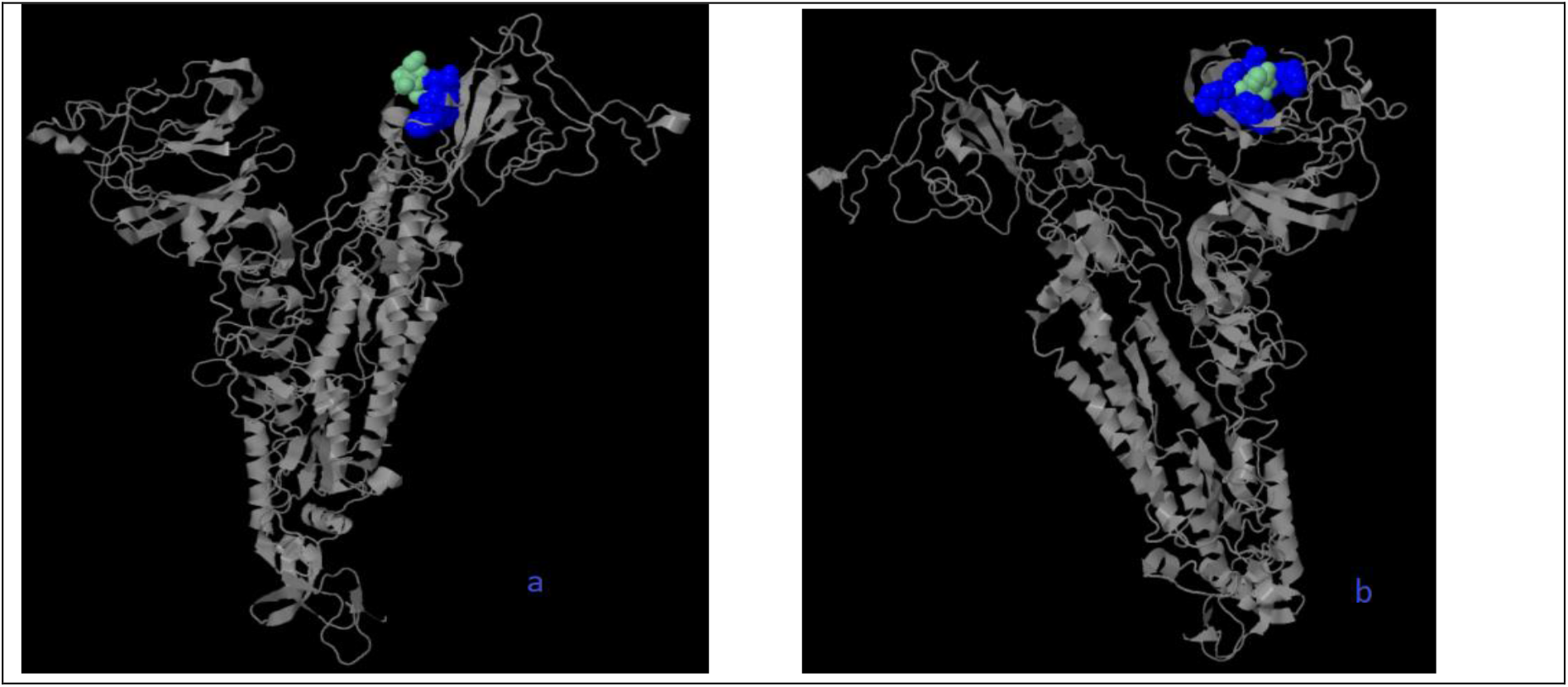
shows the ligand binding site and some varying features when two or more mutations occur. While blue color represents predicted residue, cyan represents heterogens based on 3DLigandStie method.

QJX45344 isolated from Africa has two mutations at the same sequence and the first (a on the Figure 4) represents result of sequence.. Like figure 3, source are used to predict of structure is 2dd8_S, 2ajf_E but predicting binding sites are different from figure 3. They are 338 PHE(contact:1, Av distance:0.00), 339 GLY (contact:1, Av distance:0.00), 342 PHE(contact:2, Av distance:0.18), 343 ASN (contact:2, Av distance:0.00) respectively. The second (b on the Figure 4) is about QLA46612 (has three mutation) whole sequence isolated from South Korea. Sources are used to predict of structure 1ww6_A, 1ulf_A, 1ulc_B and predicting binding site are 118 LEU (contact:3, Av distance:0.27), 120 VAL (contact:3, Av distance:0.16) 127 VAL (contact:3, Av distance:0.05), 129 LYS (contact:3, Av distance:0.00), 157 PHE (contact:3, Av distance:0.169), 159 VAL (contact:3, Av distance:0.00), 160 TYR (contact:2, Av distance:0.00), 169 GLU (contact:2, Av distance:0.54). It tend to be different from figure 3. I may say more than one mutations effect on ligand binding site based on 3DLigandStie method.

## 4. Conclusion

It is said that D614G increases infectivity of the COVID-19 Virus, like this idea other mutation predicted damage structure may increase infectivity [16]. As well as this mutation, my study reveals that there are many mutations are shown table 1–5 and some of them are seen all regions even some belongs specific region. For example D614G is seen all regions even P295H is seen only Asia. One can see all mutations regions by using access. number on the tables. In addition, more than one mutation was detected in sequences isolated in some regions specially in Asia even four mutations were seen in the same sequence. There may be human mistake, but when these four mutations were used, it was seen that the spike did not affect the structure (Figure 3). On the other hand, some of these mutations were seen to affect the ligand binding site (Figure 4).

